# Retrieval of binding sites across the AlphaFold human proteome using protein language model representations

**DOI:** 10.64898/2026.08.30.748136

**Authors:** Keshav Mohan, Yash Bhargava

## Abstract

Protein language models (PLMs) provide powerful representations of protein sequence, but their utility for proteome-scale binding-site retrieval remains unclear. Here, we present PocketScope, a training-free framework that represents cavity-lining residues using frozen ESM-C 600M embeddings and retrieves related binding sites through exhaustive late-interaction MaxSim, without pooling or approximate nearest-neighbor search. PocketScope identified 153,805 cavities across 37,682 proteins in the AlphaFold human proteome and recovered documented drug off-targets across a curated set of pharmacological pairs. On the ProSPECCTs benchmark, PocketScope ranks 1st of 23 methods by mean rank across the ten collections. PocketScope provides a practical framework for proteome-scale off-target prediction. PocketScope is open source and also freely available as a web server at https://www.bhargavaresearch.org/pocketscope.

## 1 Introduction

Recent advancements in drug discovery have allowed for the design of inhibitors with high affinity for target proteins. Specificity, however, remains an underaddressed problem in drug discovery, as small molecules often bind multiple proteins beyond the desired target. Polypharmacology can have both detrimental and beneficial consequences, with unintended target binding contributing to adverse effects while also creating opportunities to repurpose existing compounds for new therapeutic applications. Anticipating which additional proteins a drug will bind is therefore a central problem in drug discovery, and one of the dominant structural approaches to it is pocket comparison. Pocket comparison rests on the structural intuition that two proteins can share almost nothing else about their sequence and still present the same shape and chemistry at the one cavity a drug actually touches. If a binding cavity in one protein geometrically and chemically resembles a cavity in another protein that a known drug occupies, the second protein is likely a plausible off-target for that drug. Extending this comparison across an entire proteome reframes off-target prediction as a large-scale retrieval task. Two developments make proteome-scale pocket comparison feasible. AlphaFold2 supplies accurate predicted structures for the entire human proteome ^1,2^. Protein language models supply numerical descriptions of proteins directly from sequence, without alignment or hand-engineered features. Together these make it practical to compare protein structure and chemistry across a whole proteome efficiently. PocketScope is a database assembled from both, with AlphaFold2 structures supplying the cavities, a frozen protein language model supplying the residue features, and a chemical description vector supplying the chemistry alongside them.

ESM is a protein language model trained by masked-token prediction on UniRef, a protein sequence database, whose learned representations capture structural and evolutionary information despite using no explicit structural, functional, or evolutionary labels ^3^. Those representations support varianteffect prediction, and at 650M parameters they carry enough information for a folding head to be trained on top of them ^3,4^. Understanding what information protein language model representations encode is an important question in protein modeling. These properties are especially relevant when residue-level embeddings are used to represent binding sites, where the way individual residue features are combined may affect which pockets are considered structurally similar. We explored the application of these residue features for off-target prediction and compared them against existing methods.

Current computational approaches to off-target prediction fall into three families, summarised in Table 1 according to how they represent and compare potential binding sites. Ligand-centric methods compare the query drug to compounds with known targets and transfer annotations across chemical similarity, as in SwissTargetPrediction’s combined 2D/3D fingerprint search ^5^. These are fast, well validated and widely used, but a target with no chemically similar annotated ligand is invisible to them, which is exactly the case where off-target prediction is most needed.

**Table 1.**
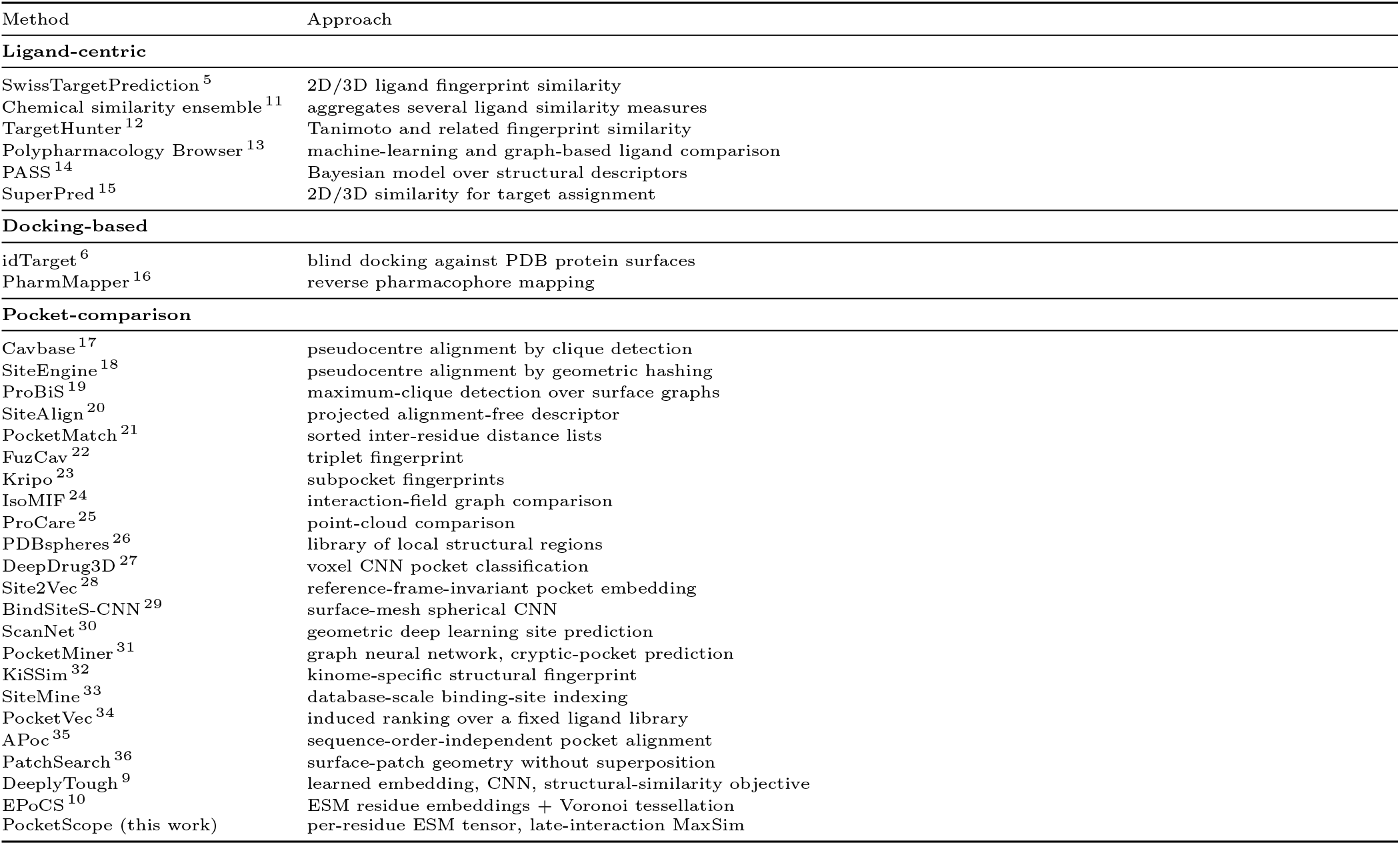
Representative methods for off-target and binding-site prediction, by family.

Docking-based target identification inverts the usual screen, docking one ligand against many structures, as id-Target does across the PDB ^6^. This uses the drug explicitly, which is an advantage in principle, but ranking many targets by docking score is expensive and the scores are known to track cavity and ligand size ^7,8^.

Pocket-comparison methods compare binding sites directly, independently of any bound ligand, and our method belongs to this family. DeeplyTough, which learns a pocket embedding with a convolutional network trained on a structural-similarity objective, is the closest learned precedent to this work, and EPoCS, which combines ESM residue embeddings with Voronoi tessellation into multiscale binding-site representations scored by Euclidean distance, is the closest in its use of a protein language model ^9,10^.

Like these methods, PocketScope compares binding sites to find off-targets, but at proteome scale. It embeds every pocket from a database of AlphaFold structures with ESM, keeps each as a per-residue tensor rather than a pooled vector, and retrieves over the whole pocketome by exact late-interaction search, previously applied to protein homolog search over per-residue embeddings ^37,38^.

## 2 Materials and methods

### 2.1 Pocket database

Structures were taken from the AlphaFold Protein Structure Database for the human proteome, UniProt reference proteome UP000005640 ^1,2^. Cavities were predicted with P2Rank 2.5.1 at default settings, using the FastRandomForest model and standard feature set, surface point sampling at a solvent radius of 1.6 Å with tessellation 2, a prediction point threshold of 0.35, a minimum cluster size of 3, a clustering distance of 3.0 Å and a neighbourhood radius of 6.0 Å ^39^.

P2Rank was run over 66,965 human AlphaFold structures, of which 37,682 proteins yielded at least one cavity, giving 153,805 pockets. We define a pocket as the set of residues lining the cavity as reported by P2Rank, truncated at 64 residues in P2Rank rank order, which gives a residue-level representation of each putative binding site for the embedding step below.

We evaluated a range of pocket-size cutoffs against a ligand-site reference built from the whole PDB and found a 10-residue cutoff to be a good operating point, removing 49.0% of predicted cavities while retaining 96.7% of ligand-site recall (Figure 2). This leaves 78,443 cavities across 28,189 proteins. The database retains all predicted cavities, so each query is scored against the full set of 153,805 cavities across 37,682 proteins, and all ranks reported here are relative to that complete set.

### 2.2 Pocket representation

Each pocket is represented as a tensor (Figure 1). For a pocket of *L* residues with one-letter sequence *s* and C*α* coordinates **x** ∈ R^*L×*3^, the per-residue match feature is

**Figure 1.**
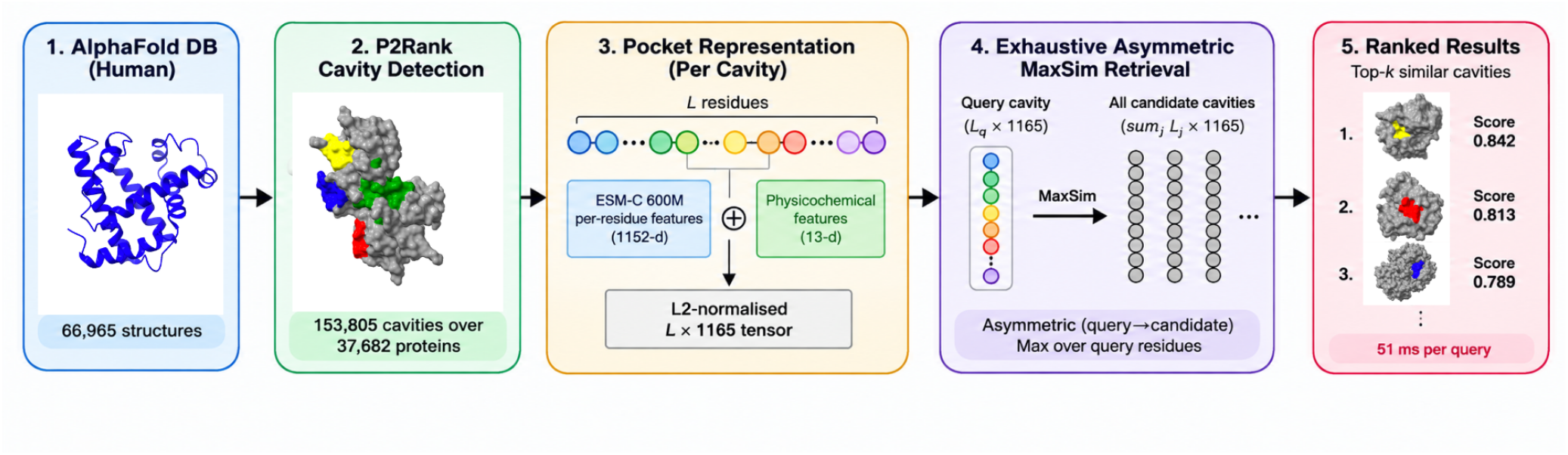
The PocketScope pipeline, from AlphaFold structures and P2Rank cavities through frozen ESM-C 600M per-residue features to exhaustive MaxSim retrieval.

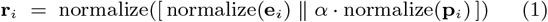

with *α* = 0.5, where **e**_*i*_ R^1152^ is the last-hidden-layer per-residue embedding from ESM-C 600M, used as released with none of its weights altered, and **p**_*i*_ R^13^ is a physicochemical descriptor. The whole chain is passed through the language model, and the vectors for the cavity-lining residues are taken from that pass and assembled into the pocket tensor. The descriptor holds seven scalars normalised to approximately [1, 1] (Kyte–Doolittle hydropathy, formal charge, side-chain volume, aromaticity, side-chain hydrogen-bond donor and acceptor counts, polarity) followed by six multi-hot pharmacophore bits (positive, negative, polar, aromatic, hydrophobic, other).

The two blocks are L2-normalised separately and then joined along the feature axis, so the 1152 language-model dimensions and the 13 chemical dimensions become one 1165-dimensional vector per residue rather than two vectors carried and compared separately. Normalising the blocks apart first lets the 13-dimensional chemical block hold a controlled share *α* of the match instead of being outnumbered by the much larger language-model vector, and the joined vector is renormalised. A pocket is therefore an *L* × 1165 tensor.

### 2.3 Retrieval by MaxSim

The input is one pocket and the output is a ranked list of candidate off-targets. The user supplies a single query cavity, either a pocket detected on a structure or a pocket of a UniProt accession already in the database, and it is turned into a query tensor by Equation 1. Every pocket in the database is then scored against it and returned in order of decreasing similarity.

Given a query tensor 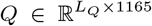, every database pocket *P* is scored by asymmetric MaxSim (Chamfer similarity). MaxSim is the late-interaction operator used in neural information retrieval, where query and target are only compared after being encoded separately ^37^,

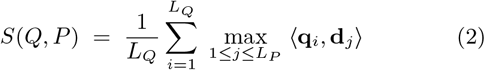

Here *L*_*Q*_ and *L*_*P*_ are the numbers of lining residues in the query cavity and in the candidate pocket *P*, so the sum runs once over each query residue and the max once over each candidate residue. Each query residue contributes its similarity to the closest matching candidate residue, and the score is the mean of those *L*_*Q*_ terms.

### 2.4 Ligand-confirmed pocket reference set

To obtain a reference set of pockets confirmed to bind a ligand, independent of P2Rank’s own prediction, we assembled accession lists for five cofactor classes (ATP, GTP, HEME, FAD, NAD), giving 1,153 accession–class entries, or 1,117 after removing 18 accessions annotated to more than one class (two entries each). For each accession, UniProt binding-site annotations were retrieved and the P2Rank pocket with maximum residue overlap (*>* 0) with the annotated site was selected, yielding 1,060 pockets (94.9%) with confirmed binding-site correspondence ^40^. We used this set to validate the 10-residue filter against an independent definition of a binding site (Section 3.1).

### 2.5 Off-target evaluation pairs

To assess whether PocketScope recovers known clinical off-targets, we evaluated its retrieval against a curated set of 30 documented pharmacological off-target pairs spanning 23 drug classes. Each pair is two proteins joined by a drug reported to bind both, with a documented clinical or pharmacological consequence; sources are listed in Supplementary Table S2. All thirty pair a protein with a paralogue, which are the cases a sequence alignment already covers.

To assess a broader space of off-target relationships, and to see how retrieval behaves as the two proteins diverge, a second and larger set was assembled from ChEMBL ^41^. Every pair of human proteins sharing a reported binder gave a pool of candidate pairs, of which 855 were retained across ten equal-width sequence-identity bands (100 pairs per band from 0–70% identity, 95, 46 and 14 in the 70–80%, 80–90% and 90–100% bands, the last two limited by how many pairs exist that are identical). Each query protein’s cavities were scored against every cavity in the database by Equation 2. A pair of proteins was scored by the highest-scoring cavity of the query against the highest-scoring cavity of the candidate. The partner’s rank among the proteins the database covers was then recorded with the query itself excluded, and recovery is a rank of 20 or better. This comparison used ESM-C 600M in chain context, the same model and setting used throughout this work. The same collections were also evaluated across five ESM model sizes and three additional model generations (Section 3.5).

Pairwise sequence identities were computed by global Needleman–Wunsch alignment, reported as identical aligned positions divided by the length of the longer sequence.

For comparison, we also ran the same queries with Fold-seek, a widely used structural alignment tool. Each query was searched against the same AlphaFold/Swiss-Prot database used by PocketScope, with an E-value threshold of 10 ^42^.

### 2.6 ProSPECCTs database

We evaluated PocketScope on ProSPECCTs, a benchmark for binding-site comparison ^43^. It comprises ten datasets, each supplying a labelled list of pocket pairs a method should or should not call similar. Performance is the percentage of should-match pairs ranked above a should-not-match pair. All ten are reported (Table 2) and three of them pair a structure with another structure of the same protein, so they measure stability rather than the ability to tell two proteins apart, and several methods reach 100% on them.

**Table 2.** The ten ProSPECCTs collections ^43^. Counts are those of the distributed pair lists, before the 10-residue minimum of Section 2.6 is applied.

| Dataset | How it is built | Relationship | Structures | Pocket pairs |
| --- | --- | --- | --- | --- |
| P1 | identical structures, different site definitions | same protein | 326 | 106,276 |
| P1.2 | identical structures, similar ligands | same protein | 45 | 2,025 |
| P2 | NMR models of one protein | same protein | 329 | 108,241 |
| P3 | mutations altering physicochemistry | point mutant | 652 | 26,860 |
| P4 | mutations preserving shape | point mutant | 652 | 26,860 |
| P5 | identity-restricted subset | different proteins, shared cofactor | 80 | 6,400 |
| P5.2 | full set | different proteins, shared cofactor | 100 | 10,000 |
| P6 | cofactors removed | unrelated proteins, shared ligand | 115 | 62 |
| P6.2 | cofactors retained | unrelated proteins, shared ligand | 115 | 62 |
| P7 | assembled from the literature | documented similar sites | 1,151 | 56,399 |

Values for the 22 methods scored on these same lists were taken from Simonovsky and Meyers ^9,43^.

PocketScope was scored on the datasets’ own pair lists with the symmetric MaxSim defined below, under ESM-C 600M in chain context as in Section 2.5. The benchmark distributes structures with their co-crystallised ligand retained. A pocket here is therefore the set of residues with any heavy atom within 6.0 Å of that ligand, rather than the P2Rank residue set used for the proteome database (Section 2.1). The ligand is then discarded, and a pair is dropped if either pocket falls below 10 residues. A pair is scored as MaxSim (Equation 2) over the two residue tensors built by Equation 1 from exactly those residues. Proteome-scale retrieval has a query and a database, so the asymmetric form of Equation 2 applies and produces every ranking reported in this work. A benchmark pair has no query and no candidate, only two pockets, so both directions are scored and averaged,

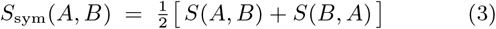

and it is this symmetric quantity that produces every ProSPECCTs number and the ablation. Averaging the directions removes any dependence on which pocket is named first, and with it part of the size dependence the asymmetric form carries.

### 2.7 Ablation analysis

To assess the contribution of pooling and the physicochemical block, we ablated each in turn, holding all other components fixed. The pooling and physicochemical-block ablations were run under ESM2-650M, before ESM-C 600M became the default encoder for the rest of this work, and the model-generation sweep below places the two side by side directly. Every ablated variant was benchmarked on all ten ProSPECCTs collections under the protocol of Section 2.6, over the same cavities and the same pair lists.

The pooled scorer mean-pools the per-residue match features of a cavity over its real residues, L2-normalises the result, and scores a pair by the cosine between the two vectors. The per-residue scorer keeps the full *L* × 1165 tensor and scores a pair by MaxSim. Pooling is the only difference between them, since both carry the physicochemical block at the same weight.

The second is the physicochemical block, removed by dropping it from the concatenation and leaving the language-model embedding alone. The third is the language model itself, varied along two axes with the cavities, the pair lists, the physicochemical block and its weight all held fixed, so that only the embedding changes. The first axis is size, running ESM2-8M, ESM2-150M, ESM2-650M, ESM2-3B and ESM2-15B, spanning 320 to 5120 dimensions per residue and a factor of 1,875 in parameters. The second is generation, running ESM-1b, ESM2-650M, ESM3-open and ESM-C 600M ^44,45^. These benchmark structures are experimental chains of a few hundred residues, so ESM-1b’s 1022-token limit binds on only 1.3% of them and no model is given a truncated input.

### 2.8 Cofactor-class classification

To ask whether the representation carries information about what a cavity binds, and not only whether two cavities match, we trained a support vector machine to tell all five cofactor classes of the ligand-confirmed set (Section 2.4) apart at once. Each pocket was averaged into a single vector, and nothing in the representation was refitted, so the classifier only reads the frozen features ^46^. Its settings were chosen inside the training data. Each class is scored against the other four, and accuracy is the area under the ROC curve built from predictions made on held-out folds.

To prevent homology between the training and test sets from inflating performance, proteins were clustered at 80% sequence identity with MMseqs2 ^47^ and whole clusters were held out together.

### 2.9 Implementation

PocketScope is available as a web server at bhargavaresearch.org*/*pocketscope and as a command-line tool, with documentation and installation instructions, at github.com*/*Bhargava-Systems-Research-Inc*/*PocketScope. The web server can also dock a compound into the top-ranked retrieved pockets using AutoDock Vina ^48^, so a candidate off-target can be followed up directly from the result list.

## 3 Results

### 3.1 Pocket database composition and the 10-residue filter

P2Rank predicted 153,805 cavities across the human proteome, strongly skewed towards small ones, with 49.0% having fewer than 10 lining residues (Figure 2, Table 3). We hypothesise that these are overwhelmingly not ligand-binding sites. To choose the threshold we built a ligand-site reference from the whole PDB rather than from a curated list. Every entry carrying a real bound ligand was taken, the ligand stripped and P2Rank run on the apo structure, which matched 119,407 of 129,495 structures and gave 2,162,492 predicted cavities, of which 532,334 touch a real bound ligand; a predicted cavity counts as recovering a ligand site when at least three of its lining residues fall within 5 Å of a ligand heavy atom. Against that reference, filtering at ten discards 49.0% of predicted cavities while preserving 96.7% of ligand-site recall (Figure 2), an enrichment of 96.7*/*51.0 = 1.90× in the usual virtual-screening sense, and leaves 78,443 cavities over 28,189 proteins. The smaller curated references agree, since only 6.4% of the 1,060 cavities confirmed against UniProt-annotated binding sites, and 4.3% of contact-level sites defined by a crystallised ligand across 1,956 experimental structures, fall below ten. Ten is also the minimum pocket length used by APoc ^35^.

**Table 3.** Composition of the pocketome and the effect of the 10-residue filter.

|  | All predicted<br>(P2Rank) | Ligand-confirmed<br>(annotation) | Ligand-defined |  |
| --- | --- | --- | --- | --- |
|  |  |  | 4 Å | 6 Å |
| Pockets | 153,805 | 1,060 | 1,956 | 1,956 |
| Proteins | 37,682 | 1,060 | — | — |
| Mean residues/pocket | 11.12 | 23.05 | 14.76 | 25.99 |
| Median residues | 10 | 21 | 14 | 27 |
| Interquartile range | 8–13 | 15–28 | 12–17 | 22–30 |
| Fraction of pockets below a residue threshold |  |  |  |  |
| < 8 residues | 24.8% | 2.0% | 0.05% | 0.0% |
| < 10 residues | 49.0% | 6.4% | 4.3% | 0.0% |
| < 12 residues | 67.7% | 12.8% | 17.0% | 0.0% |
| < 15 residues | 82.7% | 22.4% | 51.4% | 0.2% |
| Retained by the filter | 51.0% | 93.6% | 95.7% | 100% |

**Figure 2.**
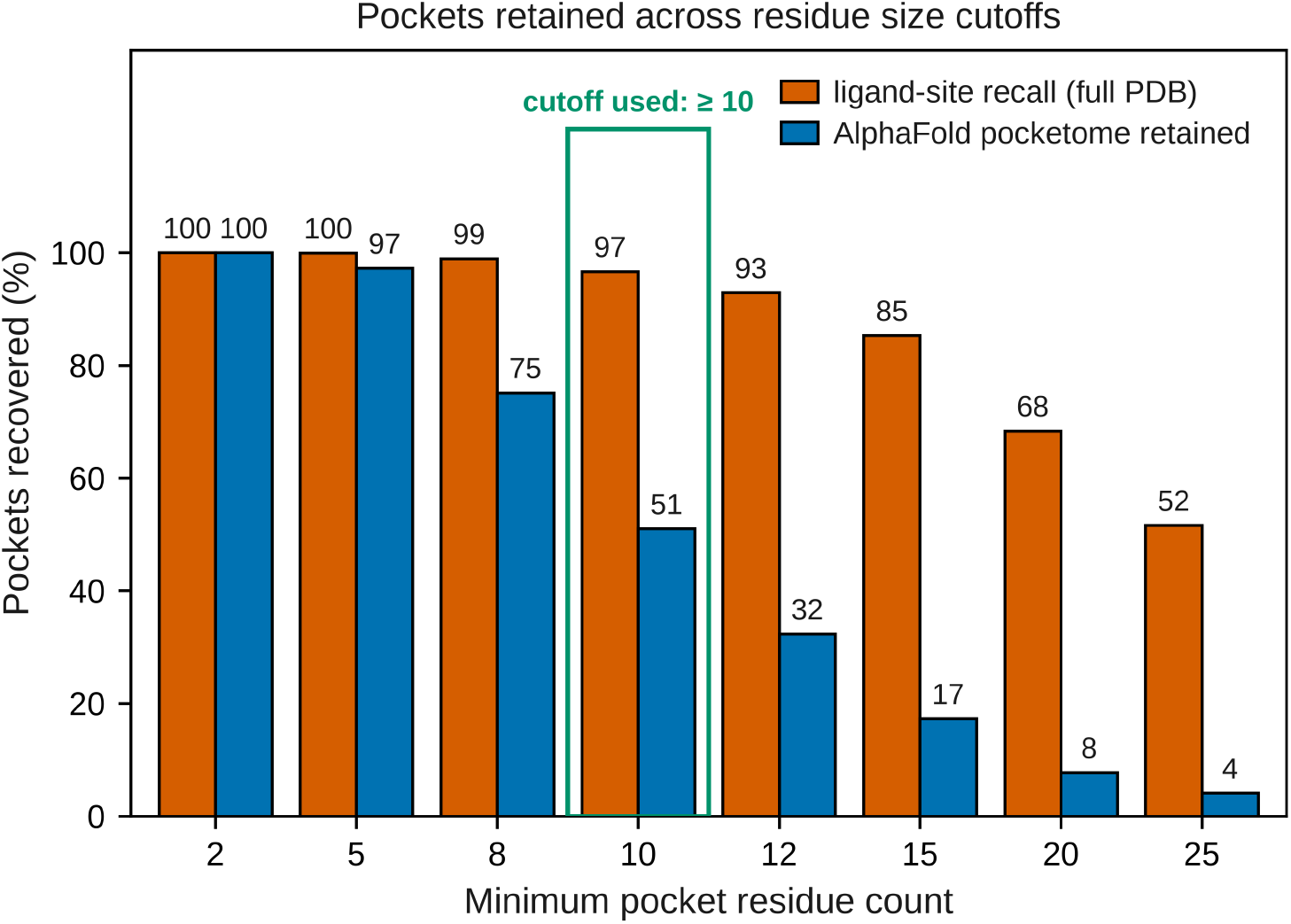
Ligand-site recall over the full PDB and AlphaFold-pocketome retention against the minimum lining-residue count.

### 3.2 Off-target retrieval on documented pairs

Queried with a primary target’s pocket, the retriever recovers documented off-target pairs across a range of target classes, spanning nuclear receptors, bromodomains, kinases, deacetylases, phosphodiesterases, transporters, GPCRs and ion-channel subunits. All 30 pairs, their ranks, the drug that joins them, its reported clinical or pharmacological consequence and the source for each are in Supplementary Table S2.

The retriever places the documented off-target inside the top 20 for 29 of the 30 pairs, ranking every protein in the database with the query protein itself excluded; the single miss is CDK2 to CDK4 at rank 25. Each of the 30 is a protein and a close paralogue, ESR1 to ESR2, ROCK1 to ROCK2, PTGS2 to PTGS1, which is where most documented off-target liability occurs, so the retrievals are pharmacologically relevant. A sequence alignment finds these pairs too; only cross-family pairs differentiate a pocket method from a more trivial alignment.

### 3.3 Comparison against Foldseek

On the 855 ChEMBL co-target pairs of Section 2.5, binned by the sequence identity of the two proteins, PocketScope (run under ESM-C 600M in chain context) matches or leads Foldseek at every identity band (Figure 3), rather than trading places with it ^42^. Both methods are near zero below 20% identity, climb steeply through the 20–40% range, and saturate by the 40–50% band; over all 855 pairs, recall at a top-20 cut is 66.1% for PocketScope against 63.7% for Foldseek.

**Figure 3.**
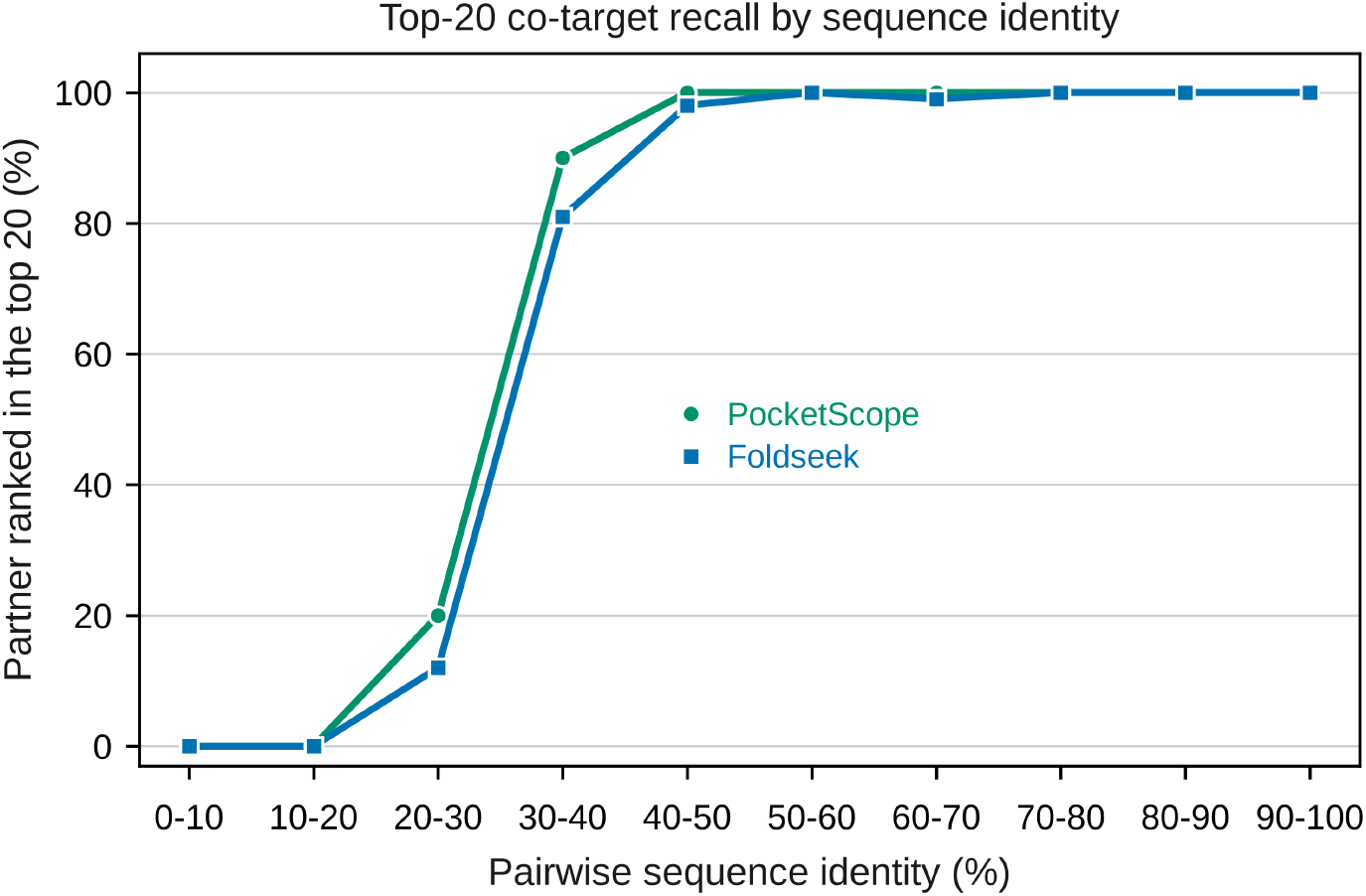
Top-20 recall of PocketScope and Foldseek on 855 ChEMBL co-target pairs, binned by pairwise sequence identity.

The two methods agree on 96.0% of the 855 pairs, retrieving the partner inside the top 20 or missing it together. They differ on 34 pairs, of which PocketScope retrieves the partner on 27 and Foldseek on 7 (exact McNemar test, *p* = 8×10^*−*4^), a difference of 2.3 percentage points with a 95% confidence interval of 1.0 to 3.7. The margin is too small to call either method the better retriever.

### 3.4 ProSPECCTs

We evaluated PocketScope against the published ProSPECCTs benchmarks to assess its ability to discriminate between related and structurally distinct protein pockets. The ten ProSPECCTs collections, the pocket definition and the scoring protocol are described in Section 2.6 and summarised in Table 2. Four of the ten pair cavities taken from distinct proteins, P5 and P5.2 on a shared cofactor and P6 and P6.2 on a shared ligand; the other six pair a structure with another structure of the same protein or with a point-mutated variant of it. Ehrt et al. scored 21 of these methods on these same pair lists and Simonovsky and Meyers added DeeplyTough (Figure 4). Ranked by mean position across the ten collections, PocketScope places 1st of 23, a mean rank of 6.1, and it is above the field median on eight of the ten collections. PocketScope reaches 100% on the three collections that pair a structure with another structure of the same protein, namely identical structures with different site definitions (P1), the same structures with similar ligands (P1.2), and NMR models of one protein (P2), where a perfect score is common. It leads the field outright on the two collections whose decoys are point mutants that alter the pocket’s physicochemistry (P3, 95.7%) or preserve its shape (P4, 94.1%), and on the documented similar sites (P7, 90.4%). In the subsets where the two proteins are genuinely different, PocketScope’s lead diminishes. It is above the field median on the full shared-cofactor set (P5.2) and on unrelated proteins sharing a ligand (P6), and below it on the identity-restricted shared-cofactor set (P5) and on the same shared-ligand pairs with their co-factors retained (P6.2), by margins too small to separate one method from another on the few pairs these collections score.

**Figure 4.**
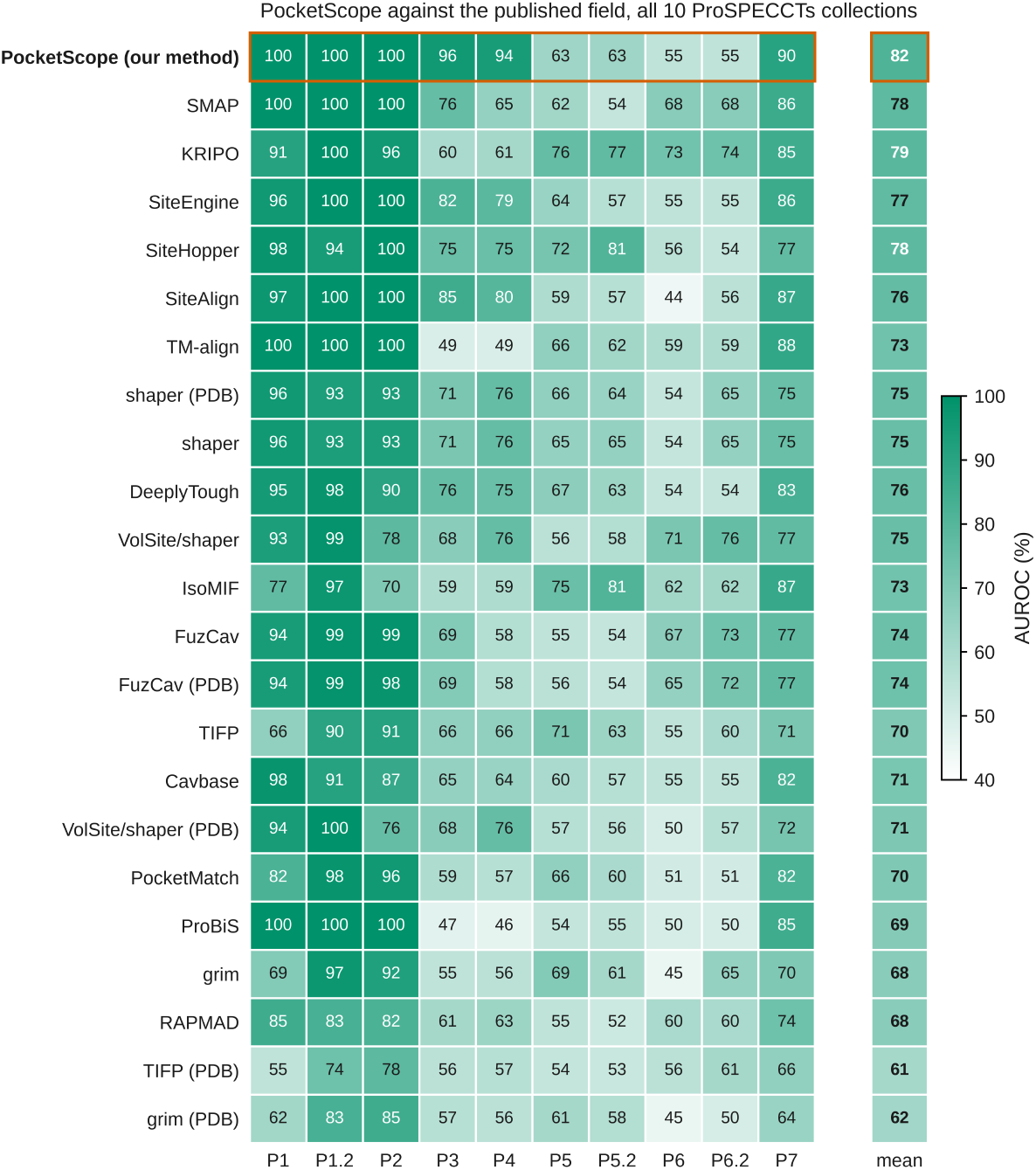
PocketScope against the 22 published methods on all ten ProSPECCTs collections. Cell values are percent of should-match pairs ranked above a should-not-match pair.

### 3.5 Representation and language-model ablations

To assess the contribution of pooling and the physicochemical block, we ablated each in turn, holding all other components fixed. The cavity is kept as a set of per-residue vectors rather than pooled into one, and a 13-dimensional physicochemical block is concatenated to each residue. We ablated both on all ten collections under the ESM2-650M default, over identical cavities and identical pair lists, so that nothing varies but the part under test (Figure 5, Supplementary Tables S3 and S4). A third ablation removes the cavity restriction itself. The whole chain is embedded as before, but every residue is kept in the tensor rather than only the cavity-lining ones, and the pair is scored by MaxSim over those full tensors (Supplementary Table S5).

**Figure 5.**
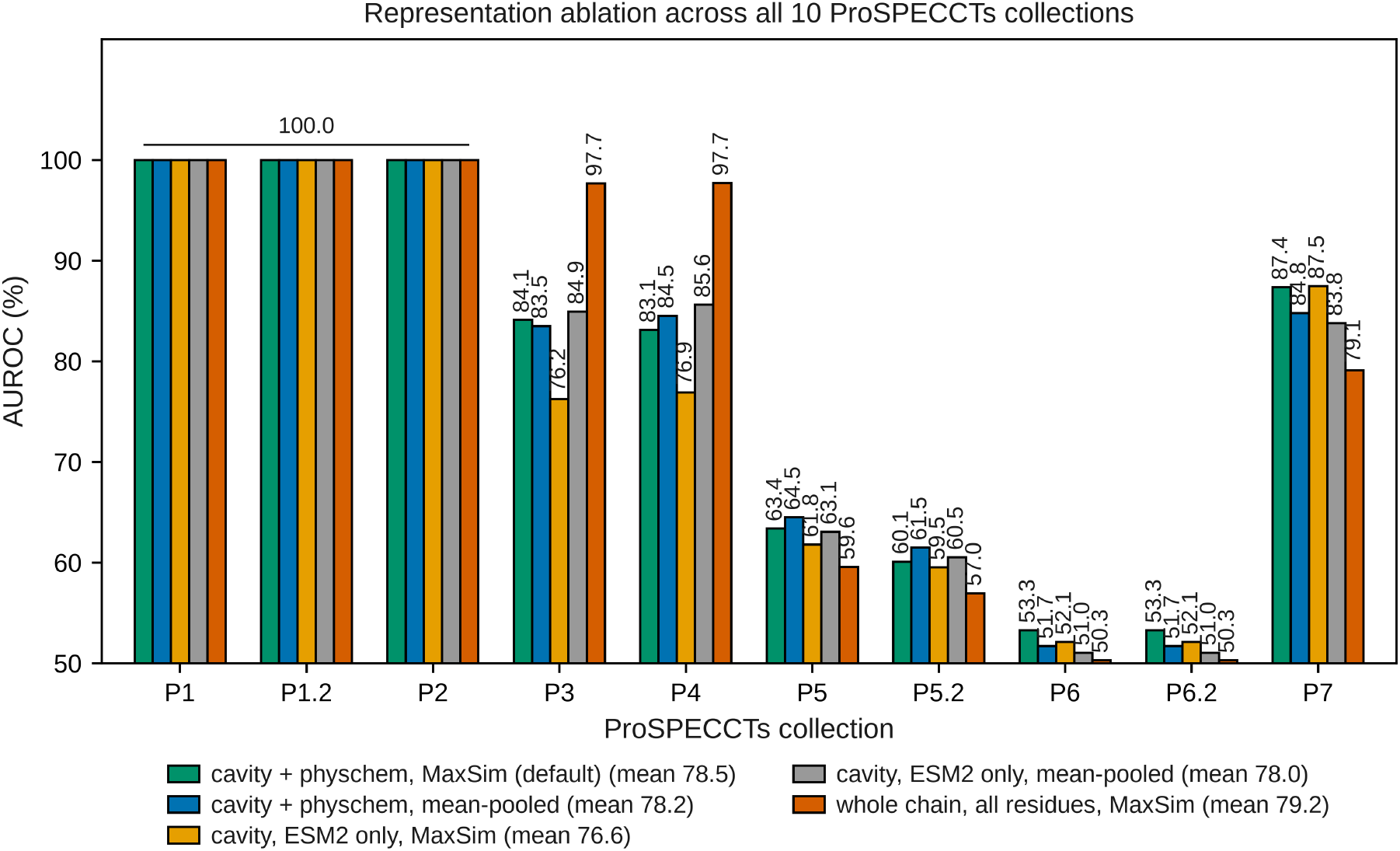
Effect of removing pooling, removing the physicochemical block, and keeping the whole chain instead of the cavity, across all ten ProSPECCTs collections.

The pooled and per-residue scorers tie at 100% on the three same-protein collections, and across the remaining seven the two track each other within a few percentage points in either direction. MaxSim enables efficient proteome-scale search without a pooling step, but on these benchmarks it buys little over pooling by itself.

The two mutated-pocket collections hold shape roughly fixed and vary chemistry, and there the chemistry-altered and shape-preserved point mutants (P3, P4) both fall by several points when the block is dropped, whether or not the cavity is pooled. On the other collections, ablating the block has minimal effect, only causing a one to two point difference, and on the documented similar sites (P7) close to nothing. This is notable as it carries no information the language model has not already seen. A plausible explanation is that the language model distributes chemical character diffusely across 1152 correlated dimensions, whereas the 13 channels state hydrophobicity, charge and hydrogen-bonding capacity explicitly and at a fixed weight, so an otherwise small signal is not diluted when the vectors are compared by cosine.

Keeping the whole chain rather than the cavity changes the answer in a way that depends entirely on what a collection asks. On the two mutated-pocket collections it rises sharply. The chemistry-altered set goes from 84.3% to 97.7%, and the shape-preserved one from 83.1% to 97.7%. A matched pair in those collections is one protein and a point-mutated copy of itself, so chain-level identity is very nearly the label.

On every collection that pairs genuinely different proteins it falls instead. The identity-restricted shared-cofactor set (P5) drops from 63.4% to 59.6%, and the documented similar sites (P7) from 86.4% to 79.1%. Unrelated proteins sharing a ligand (P6) go from 53.1% to 50.3%, a likely insignificant difference on 62 pairs. The cavity constraint prevents the score from becoming whole-protein similarity, which is something that sequence alignment can already measure.

The size of the resulting tensor rules the variant out independently of what it scores. Cavities kept at the 10-residue filter hold 1.7 million residues across the pocketome, whereas the same proteins hold 17.2 million residues in full, at a mean chain length of 456. Stored the way the index is stored, one tensor per cavity, the full-sequence version is 163 GB against the 22.9 GB of the current index, and it is 40 GB even if a single chain tensor is shared by every cavity in a protein.

Model size has no consistent effect on the representation (Figure 6, Supplementary Table S1). Across a factor of 1,875 in parameters the three same-protein collections stay saturated, at or above 99% even with the 8M model, so nothing there needs a large model at all. Size helps on the mutated-pocket collections up to 3B and then falls back at 15B, below the 650M model that is twenty-three times smaller; the one collection 15B wins outright is the documented similar sites (P7). On the collections that pair different proteins the ladder is flat and unordered, which is not a result. No model in the ladder clears chance on the unrelated-protein set (P6) by any margin that survives its confidence interval.

**Figure 6.**
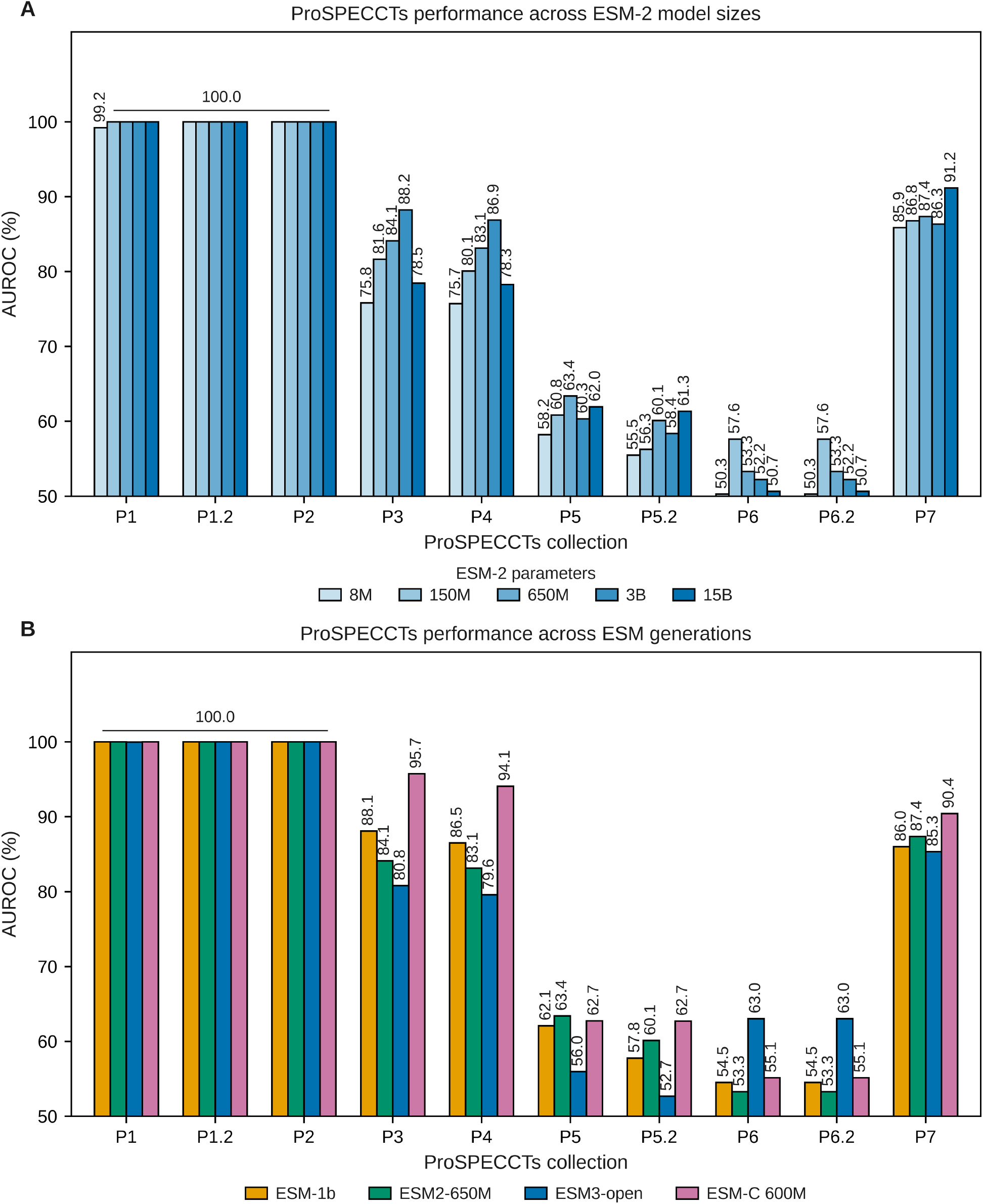
ProSPECCTs performance across ESM model sizes (A) and generations (B).

Across generations we found ESM-C 600M to be the best encoder (Figure 6B), which is why it is the one used throughout this work.

### 3.6 Cofactor classes in the representation

A single classifier separates all five cofactor classes, with no cluster of homologues spanning training and test (Figure 7, Table 4). The macro-averaged AUC is 0.991. Heme is the most distinct class at 0.999, which is unsurprising given how little its cavity has in common with a nucleotide site. The four nucleotide classes are separated almost as cleanly, between 0.983 and 0.993, even though they share an adenine-binding half. FAD is the lowest of them and also the smallest class, at 67 pockets. These embeddings are therefore rich descriptions of a binding site in their own right, carrying what the cavity binds and not only how closely it resembles another cavity.

**Table 4.** All five cofactor classes separated at once by a support vector machine on the frozen pocket representation, each class scored against the other four. AUC is the mean over 50 splits with its standard error, under 80% sequence-identity clustering.

| Class | Pockets | AUC |
| --- | --- | --- |
| ATP | 469 | $0.991 \pm 0.001$ |
| GTP | 311 | $0.991 \pm 0.001$ |
| HEME | 110 | $0.999 \pm 0.000$ |
| FAD | 67 | $0.983 \pm 0.001$ |
| NAD | 103 | $0.993 \pm 0.001$ |
| Macro average | 1,060 | $0.991 \pm 0.000$ |

**Figure 7.**
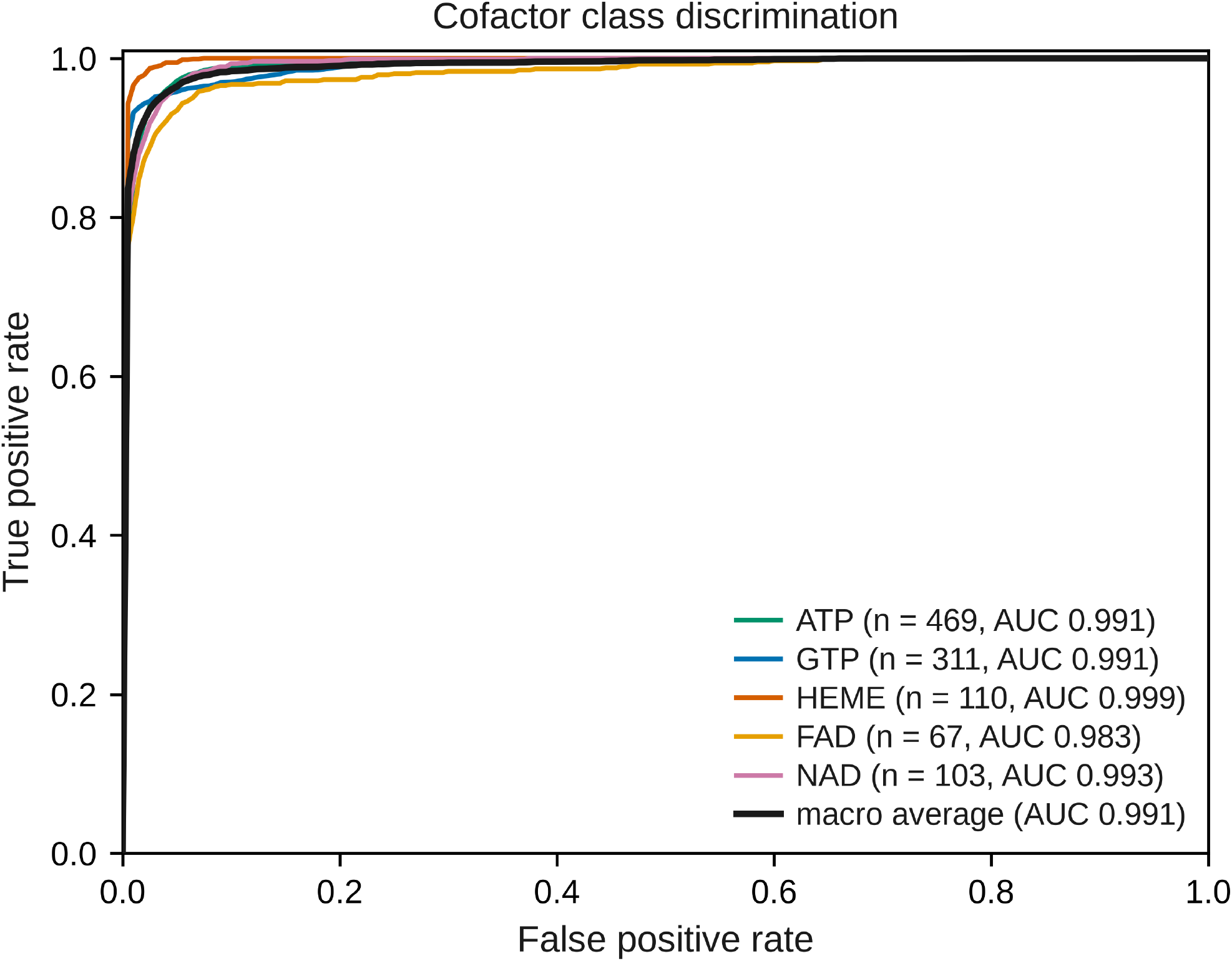
One-vs-rest ROC curves for the five cofactor classes, from predictions made on held-out folds under 80% sequence-identity clustering. Each curve is the mean over 50 splits.

## 4 Discussion

### Off-target retrieval

Queried with a drug’s target protein, PocketScope returns the documented off-target inside the top 20 for 29 of 30 curated pairs and most inside the top 5, over every predicted cavity in the human proteome and with the query protein excluded. All of them are paralogous, and that is where most documented off-target liability arises, so the retrievals are pharmacologically relevant.

### Proteome-scale search at per-residue granularity

That result comes out of an index that keeps every cavity at full resolution. We timed this directly against Foldseek on one GPU. Once each database is resident, a PocketScope query takes approximately 51 ms against approximately 2.3 s for a Foldseek search of the same AFDB/Swiss-Prot database used in Section 2.5, a roughly 45-fold difference. Of the methods compared here, PocketScope and Foldseek are the only two that search the entire human proteome for every query rather than a preselected subset of it, and PocketScope does so roughly 45 times faster. To our knowledge this is the first pocket-comparison method to search an entire human pocketome at that granularity and at interactive speed; earlier proteome-scale work either reduces each cavity to a single descriptor or precomputes a fixed ligand-ranking profile.

### ProSPECCTs benchmark

Against the 22 methods benchmarked on ProSPECCTs by Ehrt et al. and collected by Simonovsky and Meyers, PocketScope places 1st of 23 by mean rank across the ten collections, above the field median on eight of them and leading outright on three, the two mutated-pocket collections (P3, P4) and the documented similar sites (P7).

The same boundary shows up in every experiment here, and it defines the method’s operating range rather than undermining it. On the four ProSPECCTs collections that pair different proteins PocketScope splits with the field, above median on two and below on two. A binding site is defined by the three-dimensional arrangement of side chains around a void, whereas a language model encodes a property of the chain and its evolutionary context, so two unrelated proteins with a shared binder have, by construction, little chain-level similarity to find.

## 5 Conclusions

PocketScope enables large-scale binding-site searches across the proteome for potential drug off-targets. It represents 153,805 predicted human cavities using per-residue ESM-C 600M embeddings and scores each query against the full database by exact late-interaction MaxSim in about 51 ms, roughly 45 times faster per query than a Foldseek search of the same AFDB/Swiss-Prot database. Queried with a drug’s target protein, it retrieves the documented off-target among the top 20 for 29 of 30 curated pairs, with most in the top 5. Against Foldseek on 855 ChEMBL co-target pairs it matches or leads at every sequence-identity band. On ProSPECCTs, PocketScope ranks 1st among 23 methods by mean rank across the ten collections. Together, these results show that binding-site retrieval can extend off-target discovery from a small set of known relationships to a systematic search across the proteome, providing a scalable route from a drug-bound structure to candidate off-targets and experimental hypotheses.

We supply a database covering the entire human proteome, where every detected cavity carries a rich, context-dependent embedding rather than a hand-designed descriptor. These embeddings can be used downstream for clustering the human pocketome into families of chemically related sites, annotating hypothetical proteins, prioritising cavities for docking, constraining generative molecule design for selectivity, or as fixed features for models trained on small labelled sets.

## CRediT authorship contribution statement

**Keshav Mohan**: Conceptualization, Methodology, Software, Formal analysis, Investigation, Data curation, Writing: original draft, Writing: review & editing. **Yash Bhargava**: Conceptualization, Methodology, Software, Formal analysis, Investigation, Data curation, Writing: original draft, Writing: review & editing.

## Ethical approval

Not applicable. This study involved no human participants, animal subjects or patient-derived material.

## Data availability

Data supporting the findings of this study are available in Zenodo at doi.org*/*10.5281*/*zenodo.22178549. The code is available at github.com*/*Bhargava-Systems-Research-Inc*/*PocketScope.

