## Supplementary Figures and Tables for "Retrieval of binding sites across the AlphaFold human proteome using protein language model representations"

### Supplementary Tables

**Table S1.** Per-collection AUC for every language model tested, as percentages. All rows share the same cavities, pair lists and scorer, so the encoder is the only variable.

| Model | P1 | P1.2 | P2 | P3 | P4 | P5 | P5.2 | P6 | P6.2 | P7 |
|---|---|---|---|---|---|---|---|---|---|---|
| ESM2-8M | 99.2 | 100.0 | 100.0 | 75.8 | 75.7 | 58.2 | 55.5 | 50.3 | 50.3 | 85.9 |
| ESM2-150M | 100.0 | 100.0 | 100.0 | 81.6 | 80.1 | 60.8 | 56.3 | 57.6 | 57.6 | 86.8 |
| ESM2-650M | 100.0 | 100.0 | 100.0 | 84.1 | 83.1 | 63.4 | 60.1 | 53.3 | 53.3 | 87.4 |
| ESM2-3B | 100.0 | 100.0 | 100.0 | 88.2 | 86.9 | 60.3 | 58.4 | 52.2 | 52.2 | 86.3 |
| ESM2-15B | 100.0 | 100.0 | 100.0 | 78.5 | 78.3 | 62.0 | 61.3 | 50.7 | 50.7 | 91.2 |
| ESM-1b | 100.0 | 100.0 | 100.0 | 88.1 | 86.5 | 62.1 | 57.8 | 54.5 | 54.5 | 86.0 |
| ESM3-open | 100.0 | 100.0 | 100.0 | 80.8 | 79.6 | 56.0 | 52.7 | 63.0 | 63.0 | 85.3 |
| ESM-C 600M | 100.0 | 100.0 | 100.0 | 95.7 | 94.1 | 62.7 | 62.7 | 55.1 | 55.1 | 90.4 |

**Table S2.** All 30 curated pharmacological off-target pairs, with the rank at which PocketScope retrieves the off-target and the reported consequence.

| Query | Off-target | Rank | Drug or class | Reported consequence | Source |
|---|---|---|---|---|---|
| HDAC1 (Q13547) | HDAC2 (Q92769) | 1 | Pan-HDAC inhibitors | Class I cross-reactivity drives myelosuppression/QT-prolongation toxicity of pan-inhibitors like vorinostat | 10.1016/j.ejmech.2025.117998 |
| PRKCA (P17252) | PRKCB (P05771) | 1 | Pan-PKC inhibitors | Staurosporine-class pan-PKC inhibitors are non-selective across PKC isoforms; selective agents (enzastaurin) were engineered specifically to avoid this | 10.1002/jcb.23090 |
| ESR1 (P03372) | ESR2 (Q92731) | 1 | Estrogen receptor SERMs | Tamoxifen/raloxifene bind both ER subtypes; tissue-selective agonist/antagonist activity underlies SERM pharmacology | 10.7554/eLife.37161 |
| FLT1 (VEGFR1) (P17948) | KDR (VEGFR2) (P35968) | 1 | Pan-VEGFR inhibitors | Sunitinib/sorafenib inhibit all three VEGFR paralogs together as part of their multi-kinase anti-angiogenic profile | 10.1038/nbt1358 |
| KDR (VEGFR2) (P35968) | FLT4 (VEGFR3) (P35916) | 1 | Pan-VEGFR inhibitors | Sunitinib/sorafenib inhibit all three VEGFR paralogs together as part of their multi-kinase anti-angiogenic profile | 10.1038/nbt1358 |
| PDE5A (O76074) | PDE6A (P16499) | 2 | Sildenafil (PDE5 inhibitor) | Retinal off-target causes transient blue-tinted vision / photosensitivity | 10.3390/biomedicines9030291 |
| ADRB2 (P07550) | ADRB1 (P08588) | 2 | Cardioselective beta-blockers | Residual beta-2 activity of "beta-1 selective" blockers risks bronchospasm in asthmatics | 10.1183/23120541.00801-2020 |
| PTGS2 (COX-2) (P35354) | PTGS1 (COX-1) (P23219) | 2 | NSAIDs / COX-2 selective inhibitors | Drives the GI-risk (COX-1) vs cardiovascular-risk (COX-2) tradeoff; basis of rofecoxib withdrawal | 10.1186/1471-2474-8-73 |
| HDAC1 (Q13547) | HDAC3 (O15379) | 2 | Pan-HDAC inhibitors | Class I cross-reactivity drives myelosuppression/QT-prolongation toxicity of pan-inhibitors like vorinostat | 10.1016/j.ejmech.2025.117998 |
| RXRA (P19793) | RXRB (P28702) | 2 | Rexinoids (RXR agonists) | Bexarotene is a pan-RXR agonist across all RXR subtypes; basis of its use in cutaneous T-cell lymphoma | 10.1038/sj.bjc.6604320 |
| RARA (P10276) | RARB (P10826) | 2 | Pan-RAR retinoids | Isotretinoin/tretinoin act as pan-RAR agonists across RAR subtypes; basis of systemic retinoid side effects | 10.1038/sj.jid.5700418 |
| OPRM1 (P35372) | OPRK1 (P41145) | 2 | Opioids (mu/kappa cross-reactivity) | Buprenorphine is a potent kappa-receptor antagonist alongside its mu partial-agonist action; contributes to its unique pharmacology | PMID 10502307 |
| BACE1 (P56817) | BACE2 (Q9Y5Z0) | 2 | BACE inhibitors (Alzheimer's) | Verubecestat's BACE2 cross-reactivity causes documented hair/skin pigmentation changes in clinical trials | 10.1186/s13195-019-0520-1 |
| FLT1 (VEGFR1) (P17948) | FLT4 (VEGFR3) (P35916) | 2 | Pan-VEGFR inhibitors | Sunitinib/sorafenib inhibit all three VEGFR paralogs together as part of their multi-kinase anti-angiogenic profile | 10.1038/nbt1358 |

| Query | Off-target | Rank | Drug or class | Reported consequence | Source |
|---|---|---|---|---|---|
| EZH1 (Q92800) | EZH2 (Q15910) | 2 | PRC2 inhibitors (epigenetic) | Tazemetostat retains measurable EZH1 activity despite EZH2-selective design, motivating dual EZH1/2 inhibitor development | 10.6004/jadpro.2022.13.2.7 |
| ROCK1 (Q13464) | ROCK2 (O75116) | 3 | Pan-ROCK inhibitors | Fasudil is a non-selective pan-ROCK inhibitor; motivated development of ROCK2-selective agents (e.g. belumosudil) | 10.1007/s00018-009-0189-x |
| DRD2 (P14416) | DRD3 (P35462) | 3 | Antipsychotics (dopamine receptor) | Nearly all approved D2-targeting antipsychotics show poor D2/D3 selectivity contributing to side-effect profile | 10.1016/S0026-895X(25)13652-3 |
| SSTR2 (P30874) | SSTR5 (P35346) | 3 | Somatostatin analogs | Octreotide binds both SSTR2 and SSTR5 with high affinity; multi-receptor profile shapes its clinical effects | 10.1038/s41467-023-36673-z |
| ADORA1 (P30542) | ADORA2A (P29274) | 3 | Xanthines (caffeine/theophylline) | Caffeine and theophylline are non-selective antagonists across adenosine A1/A2A receptor subtypes | PMID 24859174 |
| PDE5A (O76074) | PDE6C (P51160) | 4 | Sildenafil (PDE5 inhibitor) | Retinal off-target causes transient blue-tinted vision / photosensitivity | 10.3390/biomedicines9030291 |
| BRD2 (P25440) | BRD4 (O60885) | 4 | BET inhibitors | Pan-BET inhibitors (JQ1) cross BRD2/3/4 causing dose-limiting thrombocytopenia in trials | 10.18632/oncotarget.4131 |
| PIK3CD (O00329) | PIK3CG (P48736) | 4 | PI3K-delta inhibitors | Duvelisib is a dual delta/gamma PI3K inhibitor (vs delta-selective idelalisib) reflecting shared isoform pocket structure | 10.1158/1078-0432.CCR-14-2034 |
| SLC6A2 (NET) (P23975) | SLC6A3 (DAT) (Q01959) | 4 | Monoamine transporter inhibitors | Bupropion and other NET inhibitors show measurable DAT cross-occupancy contributing to stimulant-like effects | 10.1016/j.neuroimage.2013.08.001 |
| ABCC8 (SUR1) (Q09428) | ABCC9 (SUR2) (O60706) | 4 | Sulfonylureas (KATP channel) | Glyburide's high affinity for cardiac SUR2A vs pancreatic SUR1 underlies sulfonylurea cardiovascular safety concerns | PMID 30020597 |
| PDE5A (O76074) | PDE11A (Q9HCR9) | 5 | Sildenafil (PDE5 inhibitor) | Second documented sildenafil off-target family member | 10.3390/biomedicines9030291 |
| BRD3 (Q15059) | BRD4 (O60885) | 5 | BET inhibitors | Pan-BET inhibitors (JQ1) cross BRD2/3/4 causing dose-limiting thrombocytopenia in trials | 10.18632/oncotarget.4131 |
| CDK4 (P11802) | CDK6 (Q00534) | 5 | CDK4/6 inhibitors | Palbociclib/ribociclib/abemaciclib are intentionally dual CDK4/CDK6 inhibitors due to shared ATP-pocket structure | 10.1093/jncics/pkad045 |
| CYP3A4 (P08684) | CYP3A5 (P20815) | 5 | Cytochrome P450 metabolism | CYP3A4/CYP3A5 share overlapping substrate specificity for 30-50% of clinical drugs; CYP3A5 polymorphism drives dosing variability | 10.7717/peerj.18636 |
| PDE5A (O76074) | PDE6B (P35913) | 7 | Sildenafil (PDE5 inhibitor) | Retinal off-target causes transient blue-tinted vision / photosensitivity | 10.3390/biomedicines9030291 |
| CDK2 (P24941) | CDK4 (P11802) | 25 | Pan-CDK inhibitors | Early pan-CDK inhibitors (flavopiridol/roscovitine) hit CDK2 and CDK4 together; motivated later CDK4/6-selective drug design | 10.1093/jncics/pkad045 |

**Table S3.** The pooling ablation, as percentages. Per-residue keeps the cavity as a set of residue vectors and scores by MaxSim; pooled reduces it to one vector and scores by cosine.

| Collection | Pairs are | Matched | Decoy | Per-residue | Pooled | $\Delta$ |
|---|---|---|---|---|---|---|
| P1 | same protein, different structures | 13,430 | 92,846 | 100.0 | 100.0 | +0.0 |
| P1.2 | same protein, similar ligands | 241 | 1,784 | 100.0 | 100.0 | +0.0 |
| P2 | same protein, NMR models | 7,729 | 100,512 | 100.0 | 100.0 | +0.0 |
| P3 | mutated pockets, physicochemistry altered | 13,430 | 13,430 | 84.1 | 83.5 | +0.6 |
| P4 | mutated pockets, shape preserved | 13,430 | 13,430 | 83.1 | 84.5 | -1.4 |
| P5 | different proteins, shared cofactor, identity restricted | 920 | 5,480 | 63.4 | 64.5 | -1.1 |
| P5.2 | different proteins, shared cofactor, full set | 1,281 | 8,520 | 60.1 | 61.5 | -1.4 |
| P6 | unrelated proteins, shared ligand | 19 | 40 | 53.3 | 51.7 | +1.6 |
| P6.2 | unrelated proteins, shared ligand, cofactors retained | 19 | 40 | 53.3 | 51.7 | +1.6 |
| P7 | documented similar sites | 112 | 54,992 | 87.4 | 84.8 | +2.6 |

**Table S4.** The physicochemical block ablated by sweeping its weight $\alpha$, as percentages. $\alpha = 0$ removes the block; $\alpha = 1$ gives it the weight of the full embedding.

| | Per-residue MaxSim | | | | Mean-pooled cosine | | | |
|---|---|---|---|---|---|---|---|---|
| Collection | $\alpha = 0.0$ | $\alpha = 0.25$ | $\alpha = 0.5$ | $\alpha = 1.0$ | $\alpha = 0.0$ | $\alpha = 0.25$ | $\alpha = 0.5$ | $\alpha = 1.0$ |
| P1 | 100.0 | 100.0 | 100.0 | 100.0 | 100.0 | 100.0 | 100.0 | 100.0 |
| P1.2 | 100.0 | 100.0 | 100.0 | 100.0 | 100.0 | 100.0 | 100.0 | 100.0 |
| P2 | 100.0 | 100.0 | 100.0 | 100.0 | 100.0 | 100.0 | 100.0 | 100.0 |
| P3 | 76.2 | 78.6 | 84.1 | 89.1 | 84.9 | 84.6 | 83.5 | 80.2 |
| P4 | 76.9 | 78.8 | 83.1 | 87.3 | 85.6 | 85.4 | 84.5 | 81.6 |
| P5 | 61.8 | 62.3 | 63.4 | 65.8 | 63.1 | 63.5 | 64.5 | 66.4 |
| P5.2 | 59.5 | 59.7 | 60.1 | 61.3 | 60.5 | 60.8 | 61.5 | 62.9 |
| P6 | 52.1 | 52.8 | 53.3 | 54.3 | 51.0 | 51.4 | 51.7 | 52.2 |
| P6.2 | 52.1 | 52.8 | 53.3 | 54.3 | 51.0 | 51.4 | 51.7 | 52.2 |
| P7 | 87.5 | 87.5 | 87.4 | 86.9 | 83.8 | 84.1 | 84.8 | 86.6 |

**Table S5.** Keeping the whole chain instead of the cavity, as percentages. Both columns derive from a single run over identical encoders, structures and pair lists, differing only in whether the tensor comprises the cavity-lining residues or the complete chain. Residues are the mean number retained per structure.

| | | | | Residues kept | | AUC | | |
|---|---|---|---|---|---|---|---|---|
| Collection | Pairs are | Matched | Decoy | Cavity | Chain | Cavity | Whole seq. | Δ |
| P1 | same protein, different structures | 13,430 | 92,846 | 26 | 269 | 100.0 | 100.0 | +0.0 |
| P1.2 | same protein, similar ligands | 241 | 1,784 | 26 | 298 | 100.0 | 100.0 | +0.0 |
| P2 | same protein, NMR models | 7,729 | 100,512 | 26 | 152 | 100.0 | 100.0 | +0.0 |
| P3 | mutated pockets, physicochemistry altered | 13,430 | 13,430 | 26 | 269 | 84.3 | 97.7 | +13.3 |
| P4 | mutated pockets, shape preserved | 13,430 | 13,430 | 25 | 269 | 83.1 | 97.7 | +14.6 |
| P5 | different proteins, shared cofactor, identity restricted | 920 | 5,480 | 31 | 349 | 63.4 | 59.6 | -3.8 |
| P5.2 | different proteins, shared cofactor, full set | 1,320 | 8,680 | 27 | 350 | 59.9 | 57.0 | -2.9 |
| P6 | unrelated proteins, shared ligand | 19 | 43 | 21 | 472 | 53.1 | 50.3 | -2.8 |
| P6.2 | unrelated proteins, shared ligand, cofactors retained | 19 | 43 | 21 | 472 | 53.1 | 50.3 | -2.8 |
| P7 | documented similar sites | 115 | 56,284 | 30 | 353 | 86.4 | 79.1 | -7.3 |
